# Correlated gene expression and anatomical communication support synchronized brain activity in the mouse functional connectome

**DOI:** 10.1101/167304

**Authors:** Brian D. Mills, David S. Grayson, Anandakumar Shunmugavel, Oscar Miranda-Dominguez, Eric Feczko, Eric Earl, Kim Neve, Damien A. Fair

## Abstract

Cognition and behavior depend on synchronized intrinsic brain activity which is organized into functional networks across the brain. Research has investigated how anatomical connectivity both shapes and is shaped by these networks, but not how anatomical connectivity interacts with intra-areal molecular properties to drive functional connectivity. Here, we present a novel linear model to explain functional connectivity in the mouse brain by integrating systematically obtained measurements of axonal connectivity, gene expression, and resting state functional connectivity MRI. The model suggests that functional connectivity arises from synergies between anatomical links and inter-areal similarities in gene expression. By estimating these interactions, we identify anatomical modules in which correlated gene expression and anatomical connectivity cooperatively, versus distinctly, support functional connectivity. Along with providing evidence that not all genes equally contribute to functional connectivity, this research establishes new insights regarding the biological underpinnings of coordinated brain activity measured by BOLD fMRI.

## Introduction

The brain is organized into a network of synchronized activity that has a complex and reproducible topological structure (1, 2). Resting state functional connectivity (FC) MRI, a technique which measures inter-areal correlations in spontaneous brain activity, has been particularly useful for studying functional network organization in both health and disease. Local and global features of this functional network are carefully calibrated to support healthy cognition (3) and network dysfunction is seen in numerous neurodevelopmental (4, 5) and neurodegenerative diseases (6, 7). Therefore, identifying the substrates that shape functional network organization is critical in linking molecular (e.g. gene transcription) and behavioral (e.g. psychometric) markers of disease to brain function.

Despite an abundance of prior work examining the correspondence of large-scale functional and anatomical connectivity, the precise substrates that shape functional network organization remain unknown. Modeling approaches to predict FC networks based on macro- or meso-scale anatomical connectivity networks commonly simulate mass neuronal activity by optimizing parameters that describe local population dynamics as well as the contribution of inter-areal connectivity (8–10). These approaches allow for detailed theoretical exploration regarding the relative contributions of local dynamics vs global coupling, but are limited by a lack of empirical data regarding true areal differences in function. Furthermore, analytic measures of anatomical communication appear to predict FC at comparable values (11, 12), suggesting an upper limit to the predictive validity of models based on anatomical connectivity alone.

The integration of diverse data from different scales of investigation in such models may enhance our understanding of how functional networks are shaped. Although the idea is intuitive to most that FC may be guided by a combination of factors above and beyond anatomical wiring, studies investigating how the molecular properties of a given tissue influence these functional dynamics have historically been difficult to study and remain incompletely understood. There is work emerging suggesting that gene expression and areal chemoarchitecture influence spontaneous functional brain activity. For instance, associations have been found between areal densities of excitatory receptors and strength of functional connections (13, 14). Others have found that correlated gene expression, a measure of transcriptional similarity between regions, is greater within than between functional networks, and that the genes driving these relationships are involved in ion channel activity and synaptic function (15). With that said, questions remain regarding the degree to which these relationships can be explained and might interact. Previous studies in this realm have been limited in their sparsity of regions and networks investigated and a detailed understanding of the brain’s complex network structure requires that gene expression data be comprehensively mapped onto corresponding whole-brain parcellations of structural and functional data. Furthermore, it remains unknown whether transcriptional similarities and anatomical connectivity modulate FC cooperatively, competitively, or overlap with FC uniquely depending on the connection.

Here we present a model of inter-regional FC in the mouse brain by integrating comprehensively and systematically obtained measurements of axonal connectivity (16) and gene expression data (17) from the Allen Brain Institute (ABI). We investigated whether anatomical communication capacity and correlated gene expression (CGE) contribute uniquely or cooperatively to functional network architecture. We also examined whether these relationships are homogeneously expressed across the brain or whether these dependencies change according to cortical or subcortical subdivisions. Finally, in order to examine the molecular bases of the FC signal, we examined if specific clusters of genes disproportionately support these FC patterns.

## Results

### Resting state functional connectivity of the mouse connectome

C57BL/6J mice (n = 23) were maintained under light anesthesia (1-1.5% isoflurane) and scanned in an 11.75T MRI. We computed FC (z-transformed correlations) between 160 bilateral regions of interest (ROIs) defined by Allen Mouse Brain Connectivity Atlas (AMBCA) (16) and detailed in previous work (18). ROIs excluding regions labeled as brain stem and cerebellum by the AMBCA were chosen (see methods and supplementary table 1 for a complete list of regions). Figure 1 shows qualitative clustering of the mouse functional connectome, where brain regions (nodes) are pulled together if they share strong functional connections (edges) and weak functional connections are further apart in graphical distance. Regions are colored by functional module (figure 1A) as well as anatomical assignment based on the ABI region set (figure 1B). The mouse functional connectome appears to cluster by both functional as well anatomical subdivisions; however additional variables such as polysynaptic connectivity and transcriptional similarity between regions may increase our understanding of the organization of the mouse functional connectome.

**Table 1.** Multivariate models of functional connectivity (A-J) correspond to models used in figure 2. Betas represent standardized coefficients. (G) Communicability and (M) match index are anatomical connectivity measurements. Correlated gene expression (CGE) measures gene expression similarity between regions and *G*\*CGE denotes an interaction term. Distance indicates Euclidean distance between ROI centroids and spatial adjacency (SA) is a dichotomous variable indicating whether or not two regions share a border. Full formulas for each model can be seen in supplementary figure 3.

| <u>Model</u> | <u>metric</u> | <u>beta</u> | <u>p-value</u> | <u>Model</u> | <u>metric</u> | <u>beta</u> | <u>p-value</u> |
| --- | --- | --- | --- | --- | --- | --- | --- |
| A) Structure (G)<br>$r^2 = .227$ | G | 0.4799 | $< 10^{-6}$ | G) spatial<br>topography +<br>structure<br>$r^2 = .584$ | distance | 0.45278 | $< 10^{-6}$ |
| | | | | | SA | 0.26227 | $< 10^{-6}$ |
| B) Structure<br>(G+M)<br>$r^2 = .254$ | G | 0.54327 | $< 10^{-6}$ | | G | 0.22193 | $< 10^{-6}$ |
| | M | 0.17059 | $< 10^{-6}$ | | M | 0.04621 | $< 10^{-4}$ |
| C) CGE<br>$r^2 = .412$ | CGE | 0.6421 | $< 10^{-6}$ | H) spatial<br>topography +<br>CGE<br>$r^2 = .601$ | distance | 0.38341 | $< 10^{-6}$ |
| | | | | | SA | 0.23504 | $< 10^{-6}$ |
| D) Structure +<br>CGE<br>$r^2 = .452$ | G | 0.2439 | $< 10^{-6}$ | | CGE | 0.29142 | $< 10^{-6}$ |
| | M | 0.0441 | 0.002 | I) spatial<br>topography +<br>CGE + Structure<br>$r^2 = .615$ | distance | 0.37501 | $< 10^{-6}$ |
| | CGE | 0.5231 | $< 10^{-6}$ | | SA | 0.20955 | $< 10^{-6}$ |
| E) Structure +<br>CGE*G<br>$r^2 = .488$ | G | 0.26661 | $< 10^{-6}$ | | CGE | 0.24089 | $< 10^{-6}$ |
| | M | 0.033455 | n.s. | | G | 0.14267 | $< 10^{-6}$ |
| | CGE | 0.4247 | $< 10^{-6}$ | | M | 0.00994 | n.s. |
| | G*CGE | 0.21114 | $< 10^{-6}$ | J) ALL: spatial<br>topography +<br>structure*CGE<br>$r^2 = .624$ | G | 0.16167 | $< 10^{-6}$ |
| F) Spatial<br>topography only<br>$r^2 = .549$ | distance | 0.50359 | $< 10^{-6}$ | | CGE | 0.20487 | $< 10^{-6}$ |
| | SA | 0.31871 | $< 10^{-6}$ | | M | 0.004392 | n.s. |
| | | | | | G*CGE | 0.11446 | $< 10^{-6}$ |
| | | | | | distance | 0.36544 | $< 10^{-6}$ |
| | | | | | SA | 0.18161 | $< 10^{-6}$ |

**Figure 1.**
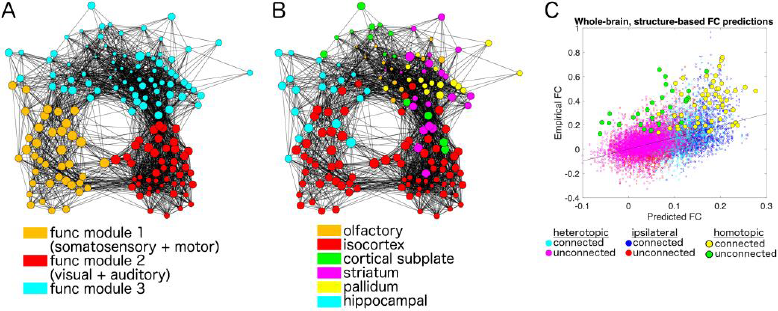
Clustering of the mouse functional connectome. Each brain region (node) shares a functional connectivity (FC) value (edge) with other nodes. Regions with strong FC are pulled more closely together and regions pairs with weak FC are moved further apart. Nodes are colored by functional (A) modularity assignment and (B) anatomical assignment. Nodes are sized by their connectivity strength (i.e., the sum of all connectivity weights to that region). C) Empirical FC values across all region pairs, illustrated as a function of anatomical connectivity network-based (i.e. structure-based) predictions of FC. Each dot represents a unique region pair, colored according to whether it is heterotopic, ipsilateral, or homotopic, and whether the region pair possesses a direct anatomical link or not. Homotopic region pairs exhibit markedly higher FC than what is predicted by structure.

### Relationships between structural and functional connectivity

Measurements of anatomical connectivity, as assessed by viral tracing, were derived from the from the AMBCA (16). Anatomical communication capacity between the 160 ROIs used in the functional analyses was then computed on the weighted structural connectivity measurements. This communication capacity metric is termed, communicability (denoted <u>*G*</u>), and is a weighted measure which describes the ease of communication between two regions (19, 20)(19, 20)(19, 20). It takes into account all possible routes between nodes (both mono and polysynaptic), but weights shorter pathways (those with fewer steps) exponentially higher. We chose this measurement of structural connectivity due to our recent work which highlights its improved capacity to model functional connectivity over simple mono-synaptic connectivity (21). We also use the matching index (denoted *M*) (22), an index which quantifies the similarity of connections between two nodes excluding their mutual connection. Matching index captures additional small contributions to FC driven by interregional similarities in connectivity patterns, as demonstrated previously (23).

Assessing the relationship between FC (unthresholded), the linear combination of G and M explained 21.7% of the variance in FC (Figure 1C) and was driven mostly by *G* (β=0.501 vs β=0.148 for *M*). Ipsilateral and heterotopic region pairs conformed closely to the overall regression line. However, homotopic region pairs showed consistently higher FC than expected by the overall regression line. This suggests an effect of functional areal similarity that is not explainable by network effects of anatomical connectivity alone.

### Inter-regional CGE, anatomical communication capacity, and spatial topology explain functional connectivity

In order to explain additional factors that contribute to the mouse functional connectome, we investigated the contribution of inter-regional CGE on resting state FC. For CGE we obtained measurements from the ABI mouse brain in-situ hybridization (ISH) data (17), which offers finely sampled whole-genome expression data within each of the allen ROIs. Due to potential differences in data quality between coronally and saggitally collected ISH data (24), we used coronally obtained genes in order to ensure the highest data quality and to avoid mixing measures from both data sets. Genetic expression of all 3188 coronally obtained ISH probes were obtained for each ROI and for each gene. Expression intensities for all genes were z-scored within each ROI and fishers z-transformed pearson correlations were computed between each anatomical region pair across genes, yielding an 80x80 matrix of inter-regional CGE. The ABI gene expression data is provided as an average of both hemispheres, thus, for all subsequent measurements G and FC were averaged across left and right hemispheres yielding comparable 80x80 matrices.

Among ipsilateral region pairs connected via monosynaptic projections, we found FC to be significantly correlated with the weighted measure of monosynaptic anatomical connectivity (*r=0.37, p=10^−6^*), but more so with communicability among monosynaptically connected ROI pairs (*G*; *r=0.435, p=10^−6^*), in accordance with previous work (21). FC and *G* were also correlated among region pairs with no monosynaptic connectivity (*r=0.334, p=10^−6^*), which is also in line with our previous work (21).

Figure 2 visualizes scatter plots for each model detailed in Table 1. The relationship between *G* and FC among all region pairs (FC is unthresholded in all models, Figure 2A/Table 1A, R^2^=.227, *β=.479*). The addition of *M* slightly but significantly increased (Figure 2B, R^2^=.254, *β=.171*) the variance explained. CGE was also strongly correlated with FC (Figure 2C, R^2^=.412, *β*=0.642). Models including the linear combination of both CGE and communicability (Figure 2D, R^2^=.452) and the addition of the interaction between CGE and G increased the variance in FC explained (Figure 2E; R^2^=.488, all individual terms except for *M*, are significant at p<10^−6^).

**Figure 2.**
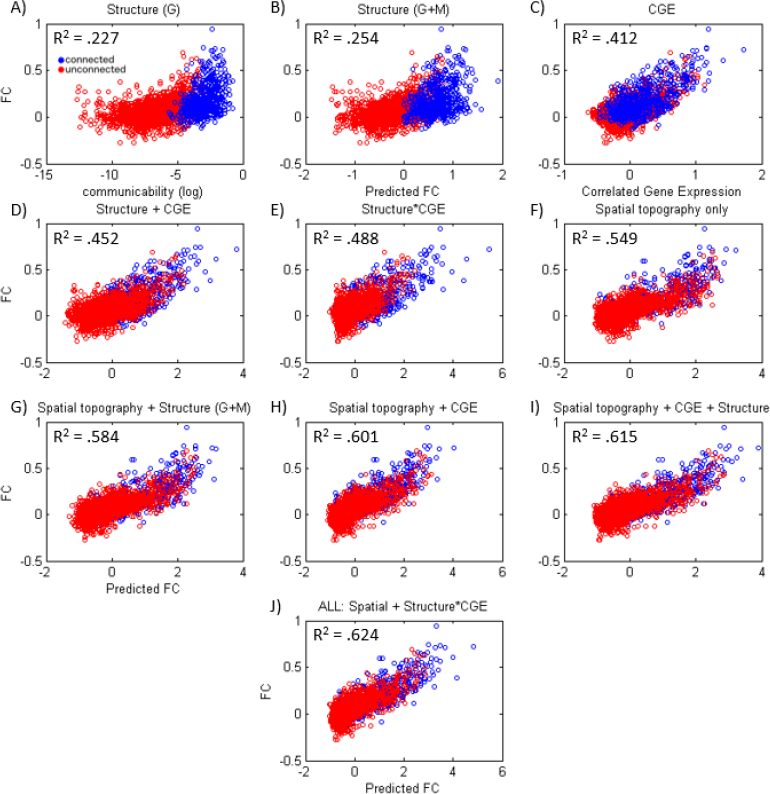
Relationships between functional connectivity (FC), anatomical connectivity measures communicability (*G*) and matching index (M), and correlated gene expression (CGE). Blue points indicate region pairs which share a direct monosynaptic anatomical connection and red points are region pairs which are anatomically unconnected. FC is illustrated as a function of A) Anatomical structure defined by only *G*, B) Anatomical structure defined by the linear regression of FC on *G* and M. C) The relationship between FC and CGE. D) FC predicted by a linear combination of structure (G+M) and CGE E) FC relationship after the inclusion the interaction between G and CGE. Next we starting with a null model predicting F_ FC as a function of spatial topography (Euclidean distance + spatial adjacency), and show that each variable adds to the variance explained above and beyond spatial topography (G-I), ending with a final J) omnibus model including all variables and the interaction between structure and CGE which explains the most variance in FC. Full equations for models depicted in A-J can be found in supplementary figure 3 and beta weights and significance for each parameter are shown in figure Table 1.

As identified by previous work (24), as well as our own (see supplementary figure 4), Euclidian distance follows a logarithmic relationship with CGE. As expected, spatial topology as defined by the - log transformed Euclidian distance between connections and spatial adjacency, a binary measure of whether two connections are touching, explain a large amount of variance in the FC signal (Figure 2F, R2=.549). Critically, the addition of each variable explains more variance than spatial topology alone, with the addition of structure (Figure 2G, R^2^=.584), CGE (Figure 2H, R^2^=.601), their linear combination (Figure 2I, R^2^=.601), and interaction (Figure 2J, R^2^=.624) explaining a total amount of variance of 62.4% of the FC matrix. Equations for each model shown in Figures 1 and 2 can be found in supplementary figure 3. Overall, we show that transcriptional similarity and network effects of axonal connectivity cooperatively, but also uniquely, support FC in a manner that goes beyond anatomical distance and spatial adjacency between regions.

**Figure 3.**
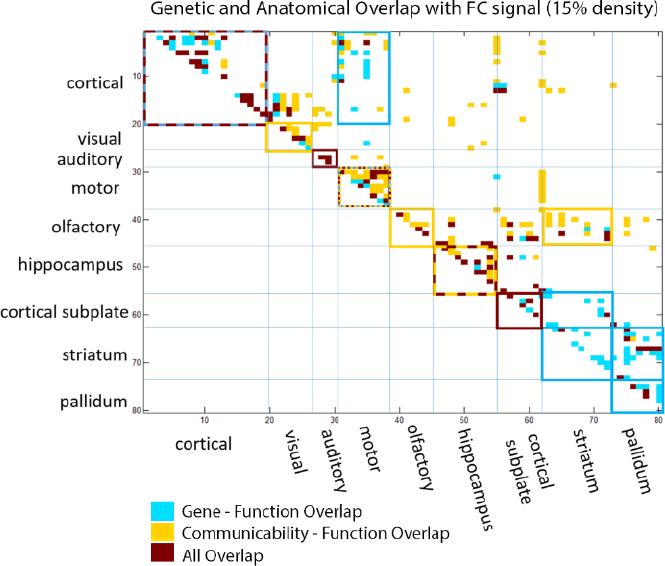
Spatial topology of genetic and anatomical overlap with the functional connectivity signal. The strongest functional connections (FC), correlated gene expression (CGE) and anatomical communicability (G) matrices were binarized (top 15% density) and overlapping patterns were compared. Connections with shared overlap, either shared between FC and CGE (aqua), FC and G (yellow), or between all three metrics (red) are shown. Anatomical modules which show significant overrepresentation of one category are outlined (based on an FDR corrected chi-squared test). Note the strong overlap between CGE and FC in the striatum and pallidum and that distant connections are primarily driven by overlap between G and FC.

**Figure 4.**
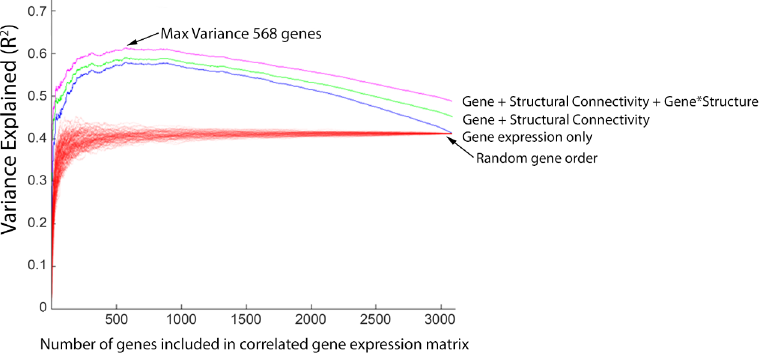
A subset of genes support the relationship between CGE and functional connectivity. Genes were rank ordered (x-axis) based on their contribution to the CGE-FC correlation (after correcting for Euclidian distance). Then, each model predicting FC was refit after incrementally adding each gene to the CGE matrix. The max variance was observed after the inclusion of 568 of the most explanatory genes to the CGE matrix, r^2^ = .6131). Each red line indicates a different permutation for gene expression only, where gene rankings were randomized on each permutation. 100 permutations are shown.

### Distinct anatomical modules are responsible for the contribution of correlated gene expression and anatomical communicability to the functional connectivity

Prior findings suggested that CGE, independent of anatomical connectivity, may help shape/explain functional connectivity networks. As such, we then explored whether this relationship is heterogeneous or homogenous across the brain. That is, are there anatomical subdivisions in which there are distinct or overlapping relationships between anatomy, gene expression, and functional connectivity? Specifically, we examined which functional connections may be supported uniquely by CGE, G, or by a combination of CGE and G. First, in order to explore these patterns we binarized each matrix at the top 15% strongest connections in the FC, CGE, and G matrices (see supplementary materials for 5%, 10%, and 20% connection densities). In order to examine overlapping profiles, a matrix was derived indicating whether there was overlap between FC and CGE, FC and G, or between all three matrices. We then calculated the statistical significance of this overlap by computing a FDR corrected chi-squared statistic for each anatomical module, which tests whether each overlap metric was more prevalent within an anatomical module than expected by chance, see methods for more details on the chi-squared approach.

Overlap between FC, G and CGE was non-uniform across the brain. As can be seen in figure 3, we found that overlap between the strongest FC and CGE within and between the striatum and pallidum, as well as within the isocortex. In contrast, we found that FC and G overlapped in hippocampal, olfactory, motor, visual areas, and between striatal and hippocampal connections. We also found overlap between all three metrics within cortical, auditory, motor, and hippocampal regions. Although these patterns are observed at the level of gross anatomical modules, a more granular view may also be informative. For instance, G, rather than CGE, may drive a subset of longer range cortical to subcortical connections. The degree of overlap depends on connection density, but importantly, the overall pattern is similar across thresholds (see supplementary materials for additional densities). Overall this suggests that FC may be both independently and cooperatively shaped by G and CGE, and that this relationship is non-uniform across the brain but depend on anatomical module.

### A subset of genes support the relationship between correlated gene expression and functional connectivity

It is likely that not all genes contribute equally to the variations in the FC signal. Next, we asked whether all genes equally contribute to the relationship between CGE and FC, and how many genes drive this relationship. To examine this we computed the FC-CGE relationship (with and without covarying Euclidean distance). Next we removing one gene and recalculated a new CGE matrix. Then, we subtracted the variance explained in the model with all genes included in the CGE matrix from the model which was calculated on the leave one out CGE matrix, and rank ordered each gene according to the magnitude of the difference in variance explained between the full and leave one out CGE matrix. Next, after rank ordering each gene by most to least related to FC, we incrementally re-introduced each gene into the CGE matrix (i.e., each time adding back one gene before computing the CGE matrix) and re-fit each model. Figure 5 shows the variance in FC explained with each model, as a function of how many genes were reintroduced into CGE matrix. The maximum amount of variance emerged after 568 genes were included in the CGE matrix (model peak without distance R^2^ = .613, with distance in model R^2^ =.726). Similar results were found when rank ordering genes without considering distance, with a max variance at 445 genes (model peak without distance, R^2^ = .671 and with distance in model, R^2^ = .702) (see supplementary results). Note the marked decline in explanatory power of the CGE after the inclusion of additional genes beyond the peak, suggesting that a subset of genes contribute disproportionately to the observed CGE-FC relationship.

These results suggest that a limited number of genes contributed to the relationship between FC and CGE. In order to examine the functions of genes that most strongly contributed to the FC signal we performed an over-representation analyses (ORA) using the software ErmineJ.Top ranked genes within the max variance peak in Figure 5 (Max variance 568 genes) were selected and compared to the background set of genes (3079 genes). This procedure identifies clusters of genes that are overrepresented within this peak, their biological and molecular processes, and cellular components. Interestingly, as opposed to some earlier work in humans **(15)**, no clusters passed statistical significance after FDR correction, potentially suggesting that these strongest related genes are equally related to several molecular and biological processes (see discussion). With that said, Table 1 shows the uncorrected results of all significant gene clusters (p<.05; peaks identified with and without covarying for Euclidian distance), which do show interesting trends that lay fodder for future study and empirical manipulation. Most notably with these findings was the over-representation of molecular processes related to voltage-gated cation channel activity, a gene cluster that was consistently over-represented regardless of distance correction.

**Table 2.**
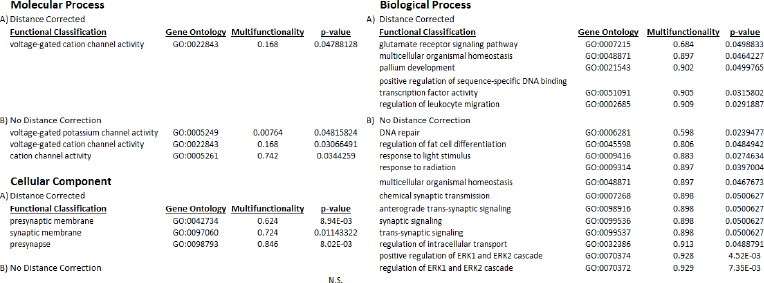
Functions of genes that support the relationship between CGE and FC. Genes which most strongly supported the CGE-FC relationship was (top genes identified in figure 5) compared the background set of all gene. Results are shown for gene rankings produced (A) with and (B) without correction for Euclidian distance between region pairs. Results are ordered by multifunctionality and were produced using the over-representation analyses in the ErmineJ software package.

## Discussion

### Modeling the influence of correlated gene expression and anatomical communicability across the functional connectome

This report investigates the fundamental neuroarchitectonics supporting synchronous large-scale brain activity. We found that functional connectivity is related to distinct aspects of structural communication (as measured via communicability) and inter-areal similarities in gene expression. Our model accounts for a significant amount of variance in the resting state functional connectivity (FC) signal (R^2^=.624) at its peak without considering improvement related to spatial proximity. The present report extends key findings from Richiardi et al., who showed in humans that correlated gene expression (CGE) is enriched within key functional brain networks relative to between these networks (15). Here we show that these principles hold in rodents and that CGE predicts functional connectivity (FC) across the brain. Interestingly, CGE explains more variance in the FC signal than anatomical communication capacity (*G*), suggesting that transcriptional similarity, and presumably similarities in protein expression, may be a crucial foundation of the FC signal in addition to anatomical wiring. We also found a significant interaction between CGE and G, suggesting that a region’s transcriptional profile and anatomical wiring may work in coordination to modulate functional synchrony with other regions. The addition of CGE to models of FC is significant given the abundance of literature that has been aimed at identifying the substrates that support functional connectivity via structural network analysis (reviewed in (25)). The present findings suggest that a more accurate modeling strategy requires the integration of structural connectivity with empirical measurements of areal molecular properties.

### Influence of anatomical connectivity and gene expression on functional connectivity is non-uniform across the brain

The contributions of anatomical communicability and gene expression to the FC signal are not spatially uniform across the brain, but cluster within and between specific subnetworks. We found that the largest overlap between the strongest CGE and FC was within non-sensory cortical areas, between cortical and somatomotor areas, within the striatal and pallidal areas, and between the striatum and cortical subplate.

Large anatomical divisions such as the striatum and neocortex display relatively uniform genetic expression compared to subcortical areas with more discrete nuclei and compared to other cortical structures such as the hippocampus that have more heterogeneous cytoarchitecture (17). The functional signal may more easily synchronize within and between areas of relatively similar physical and molecular structure, leading to increased overlap between genetic and functional metrics within the neocortex and striatum compared to the hippocampus for example.

We also observed that more distant functional connections exhibited stronger overlap with G than with CGE. Given that distant areas are more likely to be molecularly dissimilar and less likely to exhibit high CGE, these areas may rely more heavily on the presence of strong axonal pathways to facilitate functional synchrony. In the context of early development, in order to connect molecularly distinct areas across the brain, local attractive and repellent cues may guide axonal connectivity and the mapping of later functional connections between these areas.

### Relationships between physical proximity, functional and structural connectivity, and correlated gene expression

One area of caution, and interest, is the role of spatial proximity to the relationships across the three modalities. Areas that are close in spatial proximity are more likely to share more similar gene transcription profiles, have stronger anatomical connections, and share stronger functional connections. These distance relationships represent biologically meaningful information; however there is nonetheless a concern that the spatial smoothness of the fMRI signal might artifactually inflate estimates of structure-function correspondences. All fMRI data were processed without any spatial blurring in order to mitigate this possibility. Further, given the relationship between CGE and Euclidian distance, with exponentially higher CGE between regions pairs close in spatial proximity, it is critical that these type of studies properly account for spatial proximity (26). Here, we show that although highly related, these variables are also partly independent of spatial topology as anatomy and gene transcription are explanatory above and beyond null models accounting for Euclidian distance and spatial adjacency. Others have similarly corrected for spatial proximity when examining relationships between anatomy and CGE (24) or have taken alternative approaches to examine these relationships between spatially distributed functional networks (15). Given that covariation between distance and FC, G, and CGE is partly biological, we suggest that when possible the results should be compared with and without correction for spatial topology. Just as importantly, in our view, will be future experiments with a more thorough characterizing of these relationships through experimental manipulation of the expression profiles of the most strongly related genes and gene clusters – one particular advantage of conducting these experiments in rodent models.

### Correlated expression of a subset of genes disproportionately influence functional connectivity

Given that transcriptional similarity between regions had a large effect on FC, our next aim was to understand the contributions of specific gene clusters. That is, is the regional similarity across the entire transcriptome predictive, or do some genes disproportionately influence this relationship? By ranking each gene’s contribution to the CGE-FC relationship, we found that a subset of genes (568 of 3079) disproportionally drive this relationship, suggesting that covariance in particular subsets of processes may be more influential in supporting the FC signal. In this report we used gene ontology over-representation analyses (ORA) to identify the processes associated with these genes. In this section we discuss these analyses, some potential gene clusters, which may support FC, common themes, and considerations.

Although it is clear that a subset of genes disproportionately drive the relationship between CGE and FC, in contrast to previous results (15, 24), ORA on genes which are most likely to contribute to FC did not yield robust results, and as noted above, no gene cluster reached FDR corrected statistical significance. Such discrepancies here, relative to prior reports, may be due to methodological differences or differences in gene selection used for ORA. Alternatively this could suggest that across the brain a complex mixture of genes support FC and that the contribution of the genes spreads across multiple functions (at least enough such that no clusters passed correction). This could also mean that different gene clusters are critical between different anatomical connections, and that a unifying genetic function cannot describe the FC to CGE relationship across the brain. Both of these considerations deserve further investigation.

That being said, when analyzing patterns at a relaxed threshold (p < .05, uncorrected), several interesting patterns emerged yielding somewhat convergent evidence to similar reports (15, 24). For instance, genes which were most related to FC were more likely to be involved in molecular processes which were specific to voltage-gated ion channel and potassium channel activity. This is in good accordance with Richardi et al (2014) who also found preferential enrichment for these processes within compared to between functional networks. This would suggest that similar mechanisms might underlie the FC signal across species. In our report we also identified several biological processes including glutamate receptor signaling, a putative candidate for neural bases for the functional signal (27, 28), as well as cellular components related to synaptic membrane proteins and synaptic signaling.

When examining these results, it is also important to note that some gene clusters may have multiple functions that are not perfectly circumscribed to a particular process. Some genes may have many more functions than others (29) and this can be qualified by a multifunctionality score. Genes can have many diverse biological roles and highly multifunctional genes are not necessarily incorrectly assigned to a particular role, but should be interpreted with more caution (30). In our analyses, ion channels represent a gene cluster that supports FC with relatively low multifunctionality. However, we also identify additional overrepresented biological clusters with high multifunctionality scores including genes coding for homeostatic and developmental processes. These processes might be interpreted with caution as these genes also supply a rich diversity of alternative functions that, in turn, might also be related to the FC signal.

Region by region there is a complex set of neuronal, vascular, and cellular influences on the FC signal. Any given region’s activity may be modulated by neurotransmitter signaling (27, 31), the excitatory to inhibitory ratio (13, 14), local energy demands (31–33), and/or cytoarchitectural features such as synaptic density (27, 31). Each of these contributions may be reflected in interregional variability in both gene expression and spontaneous activity. Along the same lines, molecular influences on regional activity might vary across different brain subdivisions. For example, we show that the distribution of high-CGE, high-FC links are non-uniform and occurs to a greater extent in particular types of connections (i.e. striato-pallidal). Future work should address whether the gene clusters which most contribute to the functional signal vary connection to connection. Overall, the specific relationships between gene transcription and FC is far from resolved and, as noted above, will require experimental manipulation; again, highlighting the utility of preclinical animal models.

### Applications and conclusions

Models that explain functional brain organization are particularly useful in preclinical animal models, where genetic and pharmacological manipulation allow the exploration of both etiology and therapy in various neurological disorders. Structure-function relationships in the brain are known to be extremely complex, which poses fundamental challenges for neuropsychiatric research aimed at identifying substrates and treatments. In preclinical animal models and in various disorders such as autism and schizophrenia, atypical brain organization is characterized by complex and distributed disturbances of FC and structural networks (40–42). As others have noted, identifying causal links between structural and functional network disturbance is fundamentally difficult (43) and depends crucially on the validity of the measurements and the proposed model (5, 25). Here we show that models of functional connectivity can be significantly improved with the integration inter-areal similarity in gene transcription.

One of the most straightforward applications of the current statistical model would be to determine the theoretical impact of experimental perturbation to specific regions, systems, or even gene clusters within specific systems. There is evidence from work in monkeys that simulated lesions can accurately predict many widespread neurophysiological changes in response to focal empirical inactivation (21). Extending this approach to rodents, in combination with improved modeling techniques that account for transcriptional similarity, might be exceptionally useful given the time and cost associated with pharmacological screening and gene therapy testing. Recent rodent work highlights the usefulness of simulated lesions for predicting memory impairments (44), but has yet to account for the influence of areal gene expression in the modeling framework. Our results here suggest that identifying both candidate brain systems and candidate genes via simulated perturbation might be feasible for interrogating other cognitive deficits as well. Finally, experimental manipulation of activity and/or gene expression combined with simultaneous in vivo functional measurement might be used to assess what gene clusters (and what modeling framework) are best predictive of variation in typical or atypical brain function.

## Materials and Methods

### Subjects

In total, 23 C57BL/6J adult male mice ranging from 18-22g in body weight were used in the experiments. Mice were maintained on a 12-h light/dark cycle (lights on at 0600 h) at room temperature of 21 °C ± 1 °C and allowed food and water ad libitum. All experiments were performed during the animal’s light cycle. Protocols were approved by Institutional Animal Use and Care Committees of the Oregon Health & Science University and the VA Portland HCS and conducted in accordance with National Institutes of Health Principles of Laboratory Animal Care.

### Animal Preparation

Imaging in rodents generally requires the use of anesthesia to limit movement of the animals in the scanner. Here, anesthesia was induced by 3–4% isoflurane and maintained with 1–1.5% isoflurane. The selection of anesthesia may influence FC (37). Of various anesthetic regimines, we selected low dose isoflurane for the present study based on the following previous findings: 1) Functional connectivity following 1% isoflurane is preserved and comparable to that of awake mice and rats (34, 18, 35, 36, 45). 2) c-Fos activation (an immediate early gene) can be observed in isoflurane-anesthetized mice and rats (46–48). That being said, acclimated awake animals or other anesthesia regimens, such as a combination of dexmedetomidine and lower dose isoflurane (.5-.75%) (38, 39), may be an alternative.

During scanning the head was kept stationary using a custom-built head holder designed to fit in the radiofrequency (RF) coil. Respiration (80–100 bpm) and animal temperature (maintained at 37 °C) were monitored and controlled by a small animal monitoring system (Model 1030 Monitoring and Gating System; SA Instruments).

### Imaging acquisition

The imaging protocol is as described in our previous publication with slight modification (18). Imaging was performed during a single session for each animal on an 11.75T Bruker BioSpec scanner equipped with a Resonance Research, Inc. high-bandwidth shim power supply. A 20 mm ID RF quadrature volume coil (M2M, Cleveland, OH) was used for all studies. All scans were performed with Paravision 5. Using MAPSHIM, a 3D Fieldmap phase image was acquired; TR = 20 ms, TE1 = 2 ms, inter echo time = 4.003 ms, FA= 20◦, FOV= 40 mm × 18 mm × 25 mm, matrix = 80 × 90 × 125 (voxel size of 0.5 × 0.2 × 0.2 mm^3^, matching the EPI voxel size). This was followed by a T2-weighted structural image (RARE, TR = 4590 ms, effective TE = 32 ms, RARE factor = 8, 30 contiguous slices (0.5 mm thick) with interleaved acquisition, FOV= 18 × 18 mm, matrix = 150 × 150, voxel size 0.12 × 0.12 × 0.5 mm^3^, 2 repetitions). Global (volume) and local (brain voxel) shimming with MAPSHIM were performed to calculate first and second order shims prior to the functional MRI scan. The resting state fMRI consisted of a single shot gradient echo-planar imaging (EPI) sequence with the following parameters: 360 repetitions (total scan time = 15 min), TR = 2000 ms, TE = 10 ms, FA= 60◦, 30 contiguous slices (0.5 mm thick) with interleaved acquisition, FOV= 25.6 × 16 mm, matrix = 128 × 80, voxel size 0.2 × 0.2 × 0.5 mm^3^. An identical EPI sequence with 20 repetitions was acquired in the reverse phase encoding direction for topup distortion correction.

### General fMRI BOLD preprocessing

Functional images were first processed to reduce artifacts. These steps include: (1) removal of a central spike caused by MR signal offset; (2) correction of odd vs. even slice intensity differences attributable to interleaved acquisition without gaps; (3) correction of field inhomogeneities by applying topup field map correction. This required that data was collected with reversed phase-encode blips, resulting in pairs of images with distortions going in opposite directions. From these pairs the susceptibility-induced off-resonance field was estimated using a method similar to that described in (49) as implemented in FSL (50) and the two images were combined into a single corrected one. (4) movement correction; (5) within run intensity normalization to a whole brain mode value of 1000. Processed functional data was transformed to an anatomical atlas for each individual via the T2 scan. Functional data was registered to the rodent atlas supplied by the caret software (map 015 atlas) (51, 52). Each run then was resampled in atlas space on an isotropic 0.2 mm grid combining movement correction and atlas transformation in one interpolation (53).

### Rs-fcMRI pre-processing

FC pre-processed was performed as previously described with the exception of small modifications (18). Several additional preprocessing steps were used to reduce spurious variance unlikely to reflect neuronal activity (e.g. heart rate and respiration). These steps included: regression of six parameters obtained by rigid body head motion correction, the whole brain signal, and the first order derivative of the whole brain and motion parameters, followed by a temporal band-pass filter (f < 0.1 Hz).

### Regions of interest (ROIs)

160 cortical pre-defined areas (right and left hemisphere), based on the connectional and architectonic subdivisions in the mouse, as defined by the ABCA (16). ROIs within the cerebrum were used which comprise both cerebral cortical areas and cerebral nuclei. Areas defined as brain stem and cerebellum by the allen brain institute and olfactory bulb regions were not included in this analyses due to potential differences in EPI data quality. All regions included in these analyses and their anatomical module assignments (16) can be found in supplementary table 1 and their anatomical projections can be visualized on the following website (http://connectivity.brain-map.org/).

### Extraction and computation of regionwise resting state correlations

For each animal, 15 min of resting state BOLD time series data was collected. For each ROI, a resting time series was extracted post-processing and pearsons correlations were calculated for every region pair for each animal. Finally, ROI-ROI correlation, Fisher Z transformed r-values, were averaged across all subjects and used for analysis.

### Allen anatomical projection acquisition methods

Structural data were obtained from the Allen Institute for Brain Science (16). Briefly, structural data on 400 adult male C57Bl/6J mice were obtained by performing stereotaxic tracer injection (recombinant adeno-associated virus expressing EGFP anterograde tracer mapping of axonal projections), image acquisition of tracer migration (serial two-photon tomography), and data processing to make structural connection matrices. Mutual connections among 426 regions (213 ipsilateral and 213 contralateral regions) were calculated, and of these 426 regions, 168 cortical regions were used for comparison with the functional data (84 ipsilateral/right hemisphere regions and 84 contralateral/left hemisphere regions). The best fit model for connections resulted from a bounded optimization followed by a linear regression to determine connection coefficients, which assigned statistical confidence (P value) to each connection in the matrix. Structural connectivity matrices were obtained by calculating the ratio of connection density to connection strength for each ROI-ROI pair and then normalizing the ratio by the volume of the target region (ROI). For more details see Oh et al 2014.

Unlike the functional data that were undirected, the structural data contain directionality (e.g., efferent vs. afferent pathways between two nodes/ROIs). We found that this directed matrix required a very lenient threshold (P < 0.75) in order for the matrix to maintain connectedness (the ability to traverse from one node to any other node via one or more network links; a key property for making inferences regarding functional connectivity of each ROI pair). To minimize the possibility of including spurious connections, an undirected matrix was obtained by taking the average of the directed matrix with its transpose, allowing us to reduce the threshold to P < 0.05. Higher thresholds were also tested (P < 0.25) on the undirected matrix and did not alter any variance estimates by more than 1%.

Relationships between FC and anatomical connectivity were assessed using both monosynaptic connectivity strength and using a metric called communicability, which describes the ease of communication between regions via mono- and polysynaptic connections. For instance, communicability between two nodes will be stronger if there are multiple, or strong alternate paths connecting the two regions. For communicability (19) in a weighted matrix *W*, we begin by normalizing each connection weight and defining a new matrix *W’*, such that 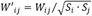, where *Si* and *Sj* are the strengths of node *i* and *j*. Communicability between *i* and *j* is defined as:

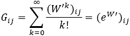

Communicability is based on the notion of communication capacity via serial relays. Given evidence that correlated activity may arise due to common efferent/afferent connectivity in the absence of communication via serial relays (23), we obtained the matching index (22) as an additional predictor of coupled activity that may not be communication-mediated. For weighted undirected networks, the matching index quantifies the similarity of connections between two nodes excluding their mutual connection, as follows where Θ(*W*_ik_) = 1 if *W*_ik_ > 0 and 0 otherwise.

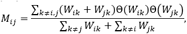

### Allen gene expression data

Gene expression data measured using in situ hybridization (ISH) from the adult C57BL/6J male mouse at age P56 were obtained from the Allen Mouse Brain Atlas (17). Expression levels of mouse in situ hybridization data from the Allen Mouse Brain Atlas were quantified using “expression energy” (fraction of stained volume * average intensity of stain), as described previously (17). Because ISH data was only available for one hemisphere, we retrieved expression energies for the same set of 80 functional ROIs used in our analyses. Because of potential differences in data quality between coronal and sagittally collected ISH data (24) we used only coronal data. To obtain this list we queried the Allen API (api.brain-map.org/api/v2/data) to obtain gene expression energy for each region in the coronal plane called using the following API query: api.brain-map.org/api/v2/data/query.json?criteria=model::StructureUnionize,rma::criteria,section_data_set[id$eqXXX],structure[id$eqYYY]. This resulted in 3318 genes, of these a final set of 3079 genes had expression energy data available for each of the 80 ROIs, which was the set used for the analyses. Gene expression energies were then normalized (z-scored) across brain regions, and pearsons correlations were computed between brain regions to assess transcriptional similarity between ROIs.

## Statistical methods

### Visualization of functional networks

Modular partitions of the network were obtained using the “Louvain” community detection algorithm (54) adapted for full, unthresholded networks with positive and negative weights (55). This algorithm identifies groups of nodes (communities, or modules) through optimization of the modularity index, or the fraction of edge weights within module partitions. The network was visualized using a force-directed graph layout where connections serve as attractive forces between nodes such that well connected groups of nodes are pulled closer together (56). In order to illustrate graphically the links between highly connected nodes, the network was thresholded at 20% (i.e. thresholded to the 20% strongest connections). Thresholds of 10% and 30% provided very similar layouts and did not alter the apparent modular organization (data not shown).

### Modeling structure-function across both hemispheres

For making FC predictions across both hemispheres, only the structural network was used since the ABI gene expression data for each region is only available as an average of both hemispheres. We employed a general linear model of FC using communicability and the matching index as the two sole predictors. FC was then plotted as a function of the predicted values from the dual-variable model. ROI pairs were plotted as separate colors depending on ipsilateral, heterotopic, or homotopic, and the presence of a monosynaptic connection.

### Modeling the transcriptional and anatomical contributions to the FC signal

Because the ABI gene expression data was only available as an average across hemispheres, the following analyses were conducted after averaging the FC networks across hemispheres as well. A series of linear regression models were assessed in order to examine the relationship between FC and transcriptional similarity, anatomical connectivity, and anatomical distance (computed as the log transformed Euclidian distance between the center of each ROI). Log transform of Euclidian distance was used in order to account for the exponentially greater functional connectivity in nearby regions. That is connectivity was better explained by the exponential than linear fit of distance on functional connectivity (24). Variance in FC explained was assessed after the inclusion of each term, as well as after the inclusion of the anatomy by CGE interaction.

### Overlap and shared and distinct connection patterns between FC, CGE, and G

In order to assess the overlap between each connection type the following analyses were conducted. First, the FC, G, and CGE matrices were thresholded and binarized, for the main analyses a 10% threshold was used. Next, for the overlap analyses, cells which were binarized for the CGE and FC, G and FC, or all three matrices were given a value of 1, and for each category, cells without overlap were given a value of 0. Similarly, for the unique and distinct analyses, matrices which were uniquely strong for CGE, G, or FC were assigned a value of 1. In order to assess the significance of these overlapping patterns we took a network level approach to see if particular anatomical clusters (defined anatomically by the ABI) were enriched for each category. This was implemented with a *χ*2 approach (see 45 for full details). Briefly, the *χ*2 test compares the observed number of binary connections (e.g., G, FC, CGE, shared, etc) within a network pair (i.e., each box figure 3) with what would be expected if the overall number of connections were evenly distributed across all network pairs. The resulting statistic is large when there are more connections than expected by chance. An empirical p-value are calculated by a permutation test, which is non-parametric and does not make assumptions about the population distribution (57, 58). Here, 10,000 permutations iterations were performed, each time randomly shuffling the binary values (i.e., CGE, G, FC, overlap, etc.) and the reported p-values for each network reflect the observed chi-square statistic compared to the permuted chi-square statistics obtained from the given network-network pair. Significant networks for each category (p<.05, FDR corrected) are outlined in Figure 3.

### Peak Analyses

In order to examine which genes were most critical for supporting the relationship between FC and CGE the following analyses was performed. First, we computed the FC-CGE correlation after including each of the 3079 genes in the CGE matrix, we then incrementally removed one gene before calculating the CGE matrix, and re-computed the FC-CGE correlation. Then, we subtracted the FC-CGE correlation from the FC-CGE correlation with one gene removed. Next, we rank ordered each gene according to how much the relationship dropped after the removal of the gene. Finally, after rank ordering each gene we incrementally re-introduced each gene (in rank order) into the CGE matrix and re-fit of the statistical models testing the relationship between FC and each predictor. We then identified the number of genes included in the CGE matrix that resulted in the highest amount of variance explained.

### Over-representation analysis

ErmineJ software, version 3.0.2 (59) was used to for over-representation analyses (ORA) comparing our target gene set corresponding to genes most related to FC (Peaks with and without covarying Euclidian distance in the gene rank list) to the background list of all coronal genes (3079 genes). Gene annotations were assigned from GO (60) using an annotation file from GEMMA (61): Generic_mouse_ ncbiIds_noParents.an was downloaded from http://www.chibi.ubc.ca/microannots/ on December 6, 2016. From the 3079 genes in our set the annotations matched 3076 genes, the final list of genes included in our ORA analyses. Over-represented biological processes, molecular processes, and cellular components were tested. We used a maximum and minimum gene set size of 100 and 20 genes, respectively, used the best scoring replicate, and for scoring we weighted each gene within the peak as 1 and the remaining background genes as -1.

## Supplementary Information

**Figure S1.**
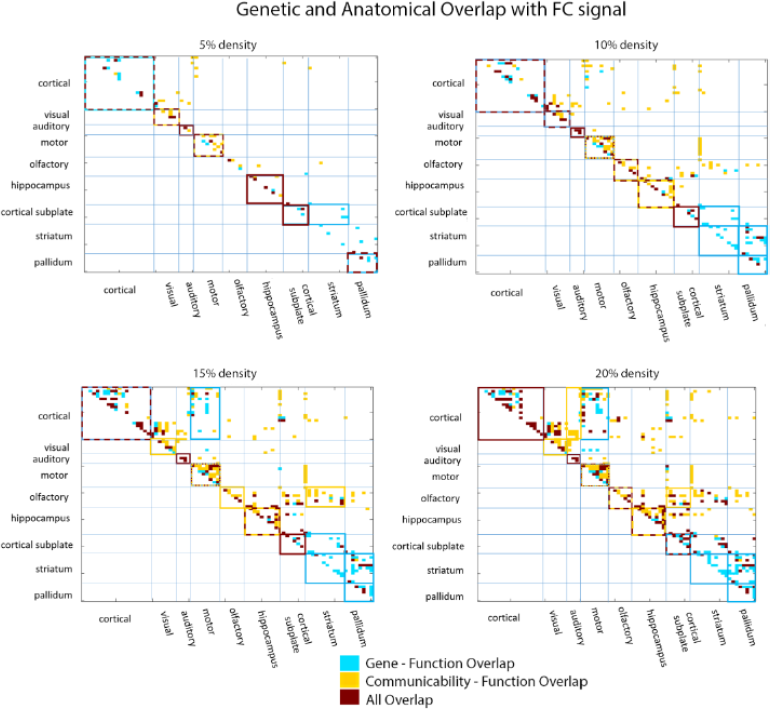
Spatial topology of genetic and anatomical overlap with the functional connectivity signal across multiple threshold densities (top 5, 10, 15, and 20%). The strongest functional connections (FC), correlated gene expression (CGE) and anatomical communicability (G) matrices were binarized at each density and overlapping patterns were compared. Connections with shared overlap, either shared between FC and CGE (aqua), FC and G (yellow), or between all three metrics (red) are shown. Anatomical modules which show significant overrepresentation of one category are outlined (based on an FDR corrected chi-squared test).

**Figure S2.**
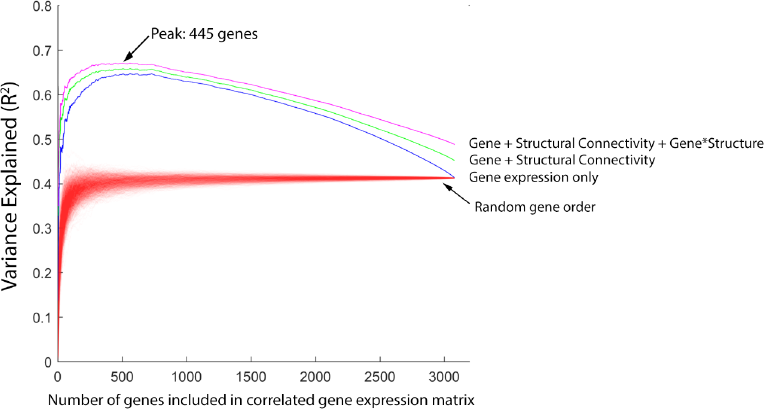
Incremental inclusion of genes into CGE matrix without distance correction during rank ordering of gene importance. A subset of genes support the relationship between CGE and functional connectivity. Genes were rank ordered (x-axis) based on their contribution to the CGE-FC correlation (after covarying Euclidian distance). Then, each model predicting FC was refit after incrementally adding each gene to the CGE matrix. The max variance was observed after the inclusion of 445 of the most explanatory genes to the CGE matrix, r^2^ = .702). Each red line indicates a different permutation for gene expression only, where gene rankings were randomized on each permutation. 1000 permutations are shown.

**Figure S3.**
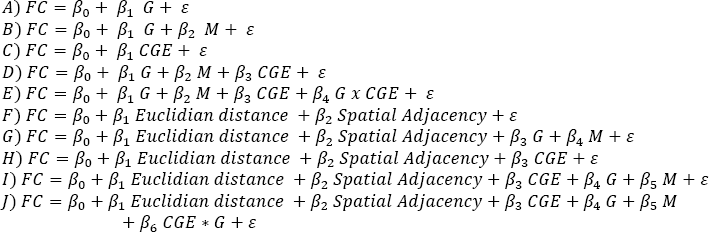
Models used to assess the relationships between functional connectivity *(FC)*, and measures of anatomical structure, communicability *(G)* and matching index *(M)*, correlated gene expression *(CGE)*, Euclidian distance between region pairs, and spatial adjacency, a binary measure of whether two regions are touching. Models A-J correspond to models used in scatters plots on Figure 2 and models described in Table 1.

**Figure S4.**
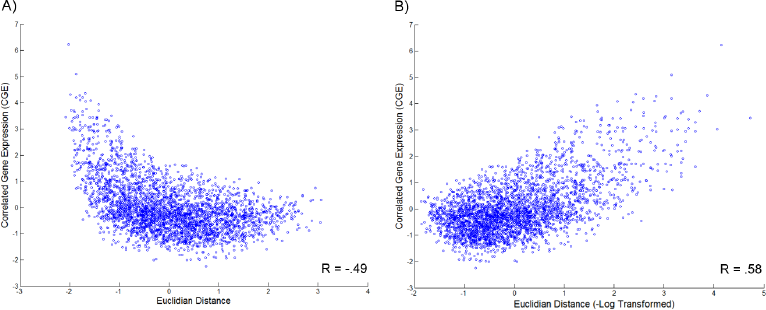
The relationship between Euclidian distance and correlated gene expression (CGE) is exponential, therefore, Euclidian distance was –log transformed before applied to all models throughout the manuscript. A) CGE by Euclidian distance (without transforming the distance matrix) shows an exponential relationship. B) The –log of Euclidean distance better fits the relationship between distance and CGE.

